# Atypical neurogenesis in induced pluripotent stem cell (iPSC) from autistic individuals

**DOI:** 10.1101/349415

**Authors:** Dwaipayan Adhya, Vivek Swarup, Roland Nagy, Lucia Dutan, Carole Shum, Eva P. Valencia-Alarcón, Kamila Maria Jozwik, Maria Andreina Mendez, Jamie Horder, Eva Loth, Paulina Nowosiad, Irene Lee, David Skuse, Frances A. Flinter, Declan Murphy, Grainne McAlonan, Daniel H. Geschwind, Jack Price, Jason Carroll, Deepak P. Srivastava, Simon Baron-Cohen

**Author notes:** Joint senior authors (Deepak P. Srivastava, Simon Baron-Cohen). Corresponding author: Deepak P. Srivastava, or Simon Baron-Cohen. Joint first authors (Dwaipayan Adhya, Vivek Swarup).

## Abstract

**Background:** Autism is a heterogenous collection of disorders with a complex molecular underpinning. Evidence from *post-mortem brain* studies using adult brains have indicated that early prenatal development may be altered in autism. Induced pluripotent stem cells (iPSCs) generated from autistic individuals with macrocephaly also indicate prenatal development as a critical period for this condition. But little is known about early altered cellular events during prenatal stages in autism.

**Methods:** IPSCs were generated from 9 unrelated autistic individuals without macrocephaly and with heterogeneous genetic backgrounds, and 6 typically developing, control, individuals. IPSCs were differentiated towards either cortical or midbrain fates. Gene expression and high throughput cellular phenotyping was used to characterise iPSCs at different stage of differentiation.

**Results:** A subset of autism-iPSC cortical neurons were RNA-sequenced to reveal autism-specific signatures similar to *post-mortem brain* studies, indicating a potential common biological mechanism. Autism-iPSCs differentiated towards a cortical fate displayed impairments in the ability to self-form into neural rosettes. In addition, autism-iPSCs demonstrated significant differences in rate of cell type assignment of cortical precursors, and dorsal and ventral forebrain precursors. These cellular phenotypes occurred in the absence of alterations in cell proliferation during cortical differentiation, differing from previous studies. Acquisition of cell fate during midbrain differentiation was not different between control- and autism-iPSCs.

**Conclusions:** Taken together, our data indicate that autism-iPSCs diverge from control-iPSCs at a cellular level during early stage of neurodevelopment. This suggests that unique developmental differences associated with autism may be established at early prenatal stages.

## Introduction

Autism spectrum conditions (henceforth autism) are a genetically heterogeneous spectrum of neurodevelopmental conditions^1–3^. Autism is characterised by impairments in social-communicative behaviours as well as repetitive behaviours. Symptoms of autism cannot be detected until twelve to eighteen months of age^4^. However, there is debate surrounding the origins of autistic symptoms. It is now well recognised that genetic factors play a key role in the emergence of autism^1, 2^. Increasing evidence indicate that perturbation during critical periods of development maybe key for the emergence of autism^5^. Moreover, autism *post-mortem brain* studies have identified dysregulation of putative prenatal gene expression pathways^6^. This suggests that early prenatal development may be a critical period for the emergence of cellular pathophysiology associated with autism^6^.

The use of induced pluripotent stem cell (iPSC) differentiated into neurons of distinct lineages^7–11^, has made it possible to study prenatal cellular behaviour in autism in detail. As iPSC-neurons contain the same genetic information as the individuals from whom they were derived, their cellular behaviours are influenced by their genetic background. Using these methods, studies have shown significant anomalies in cellular/molecular behaviour during prenatal-equivalent periods of development in autistic individuals with a co-diagnosis of macrocephaly^12–14^. These studies have demonstrated: (1) atypical neural differentiation of iPSCs fated towards a cortical lineage, and (2) an imbalance in excitatory (glutamate-producing) and inhibitory (GABA-producing) receptor activity^12, 13^. More recently, using the same collection of iPSCs, an acceleration in neuronal maturation was found to be dependent on early cortical neural precursor (NPC) development, as circumventing this stage of cortical development by direct conversion of iPSCs into mature neurons did not recapitulate these effects^14^. These phenotypes were paralleled by alterations in gene expression network dynamics during early stages of development in these iPSCs^14^. These studies highlight that the cellular and molecular phenotypes associated with autism may start before birth, and possibly at a very early stage of brain development^14^. A critical aspect of previous studies was that atypical neural differentiation observed in the autism-iPSCs was associated with higher cell proliferation^12–14^. As the autistic participants in these studies also had macrocephaly it is yet to be determined whether the observed abnormal development was due to this comorbidity. Moreover, as macrocephaly is present only in a subset of autistic individuals and it is not known if abnormal development can be generalised to autistic individuals without macrocephaly. In such individuals, it is unknown whether the acquisition of cortical fate is also accompanied by a difference in precursor population. Finally, as the majority of studies have predominantly focused on the development of forebrain/cortical neurons, it is yet to be tested whether abnormal development can also be observed in neural precursors fated towards a different lineage.

In this study, we have generated iPSCs from autistic individuals without macrocephaly and with heterogeneous genetic backgrounds to represent the wider autistic population. Initial RNA-sequencing studies using a subset of iPSC lines was used to confirm whether the transcriptomic signature of iPSC-derived neurons correlated to autistic *post-mortem* gene expression patterns. To further investigate the source of atypical gene expression, we recruited individuals from 3 independent patient cohorts to capture a wider population of autistic individuals, and undertook extensive cellular phenotyping experiments. The goal of this study was to understand if there was a fundamental difference between typical and autistic prenatal neurodevelopment, focussing primarily on early neuroectodermal structures and cell types that constitute the developing cerebral cortex.

## Materials and Methods

### Induced pluripotent stem cells

Participants were recruited and methods carried out in accordance to the ‘Patient iPSCs for Neurodevelopmental Disorders (PiNDs) study’ (REC No 13/LO/1218). Informed consent was obtained from all subjects for participation in the PiNDs study. Ethical approval for the PiNDs study was provided by the NHS Research Ethics Committee at the South London and Maudsley (SLaM) NHS R&D Office. Autistic participants were selected based on ADOS, ADI-R scores, while typical controls were selected from the population if they did not have a diagnosis of any psychiatric condition^15^. IPSC lines from autism: 9, control: 6 participants were generated from hair keratinocytes as previously described^16, 17^. Details on all participants can be found in **Supplementary Tables S1, S2, S3**). Two clones per iPSCs were used in all experiments; pluripotency of all iPSCs was determined by immunocytochemistry (**Supplementary Table S4 and Supplementary Figure 1**).

### Neuronal differentiation

IPSC lines were differentiated cortical neurons using a dual SMAD inhibition protocol which recapitulates of key hallmarks of corticogenesis^10, 17^. IPSCs were differentiated to midbrain floorplate precursors using established protocols^7, 8^. Further information can be found in Supplemental Information.

### RNA-sequencing

RNA-sequencing was performed from a subset of our cohort, on 2 clones from each participant (ASDM1, 004ASM, 010ASM, CTRM1, CTRM2, CTRM3), and each clone had 2 technical replicates. Starting with 500ng of total RNA, poly(A) containing mRNA was purified and libraries were prepared using TruSeq Stranded mRNA kit (Illumina). Unstranded libraries with a mean fragment size of 150bp (range 100-300bp) were constructed, and underwent 50bp single ended sequencing on an Illumina HiSeq 2500 machine. Bioinformatics analysis was performed using C++ and R based programs (see Supplemental Information).

### Immunocytochemistry

Differentiated iPSCs were fixed in 4% paraformaldehyde at indicated ages and processed as previously described^17^. Briefly, fixed cells were permeabilized in 0.1% Triton-X-100/PBS, and blocked in 4% normal goat serum in PBS. Primary antibodies (**Supplementary Table S5)** were incubated overnight at 4°C. Nuclei were identified by staining with DAPI. High content screening (HCS) was performed using an Opera Phenix High-Content Screening System (Perkin Elmer). Cell type analysis was performed using the Harmony Software^17^. For Rate of Cell Type Assignment (deltaCTA), the percent positively stained cells appearing per day was estimated, which was then adjusted to the total number of positive cells appearing per day in one well of a 96-well plate, assuming each well had an average of 10^5^ cells.

### Statistics

Quantification of cell types was performed using the Harmony High Content Imaging and Analysis Software (Perkin Elmer). Percentage of cells positive for desired marker versus total number of live cells was calculated. To take into consideration variability associated with iPSC differentiation, 8 independent experimental replicates of 2 clones per individual was used at every stage. Immunofluorescence was measured only from known intracellular location of markers (e.g. nucleus or cytoplasm). Independent 2-group t-test was used to check significant difference between autism and control using p-value ≤ 0.05. One way ANOVA was performed to investigate in-group variance. All statistical analysis was performed on R statistical software.

## Results

### Participant overview

Participants were recruited from the Longitudinal European Autism Project (LEAP)^18^, Brain and Body Genetic Recourse Exchange (BBGRE) studies^19^, or the Social Communication Disorders Clinic at Great Ormond Street Institute of Child Health (GOS-ICH). Of the autistic participants, eight were male and one was female (**Supplementary Table S1**). The four participants from the LEAP cohort were diagnosed with non-syndromic, while participants from BBRGE and GOS-ICH cohorts were diagnosed with syndromic autism (**Supplementary Table S2**). Syndromic participants from GOS-ICI had deletions type CNVs in the 1p21.3 and 8q21.12 regions, with *DYPD* and *PTBP2* and the *AXL* genes of note in each region respectively. Of the syndromic participants from BBGRE, two syndromic participants had deletion type CNVs in the 2p16.3 region (*NRXN1*), while the third carried a duplication in the 3p chromosomal region^20^ (**Supplementary Table S3**).

### Neurodevelopmental gene expression signatures in autism-iPSC-derived neurons

Studies using iPSCs from autistic individuals with macrocephaly have suggested an association between atypical cortical differentiation with altered cell proliferation^12, 13, 21^. We were also interested in examining cortical differentiation in iPSCs derived from individuals diagnosed with autism but without macrocephaly. Thus, we differentiated iPSCs towards a cortical fate and focused on three distinct developmental stages (**Figure 1A**): (1) Day 9: early neural precursor stage, when stem cells form new precursor cells which self-organise into neural tube-like structures known as neural rosettes with a directional apical-basal arrangement; (2) Day 21: late neural precursor stage, a period during which neural progenitor cells begin forming layers from the apical surface and are primed for differentiation into neurons as they move outwards; and (3) Day 35: immature cortical neurons, a stage at which precursors become post-mitotic and adopt a deep layer neuronal identity (**Figure 1B)**.

**Figure 1:**
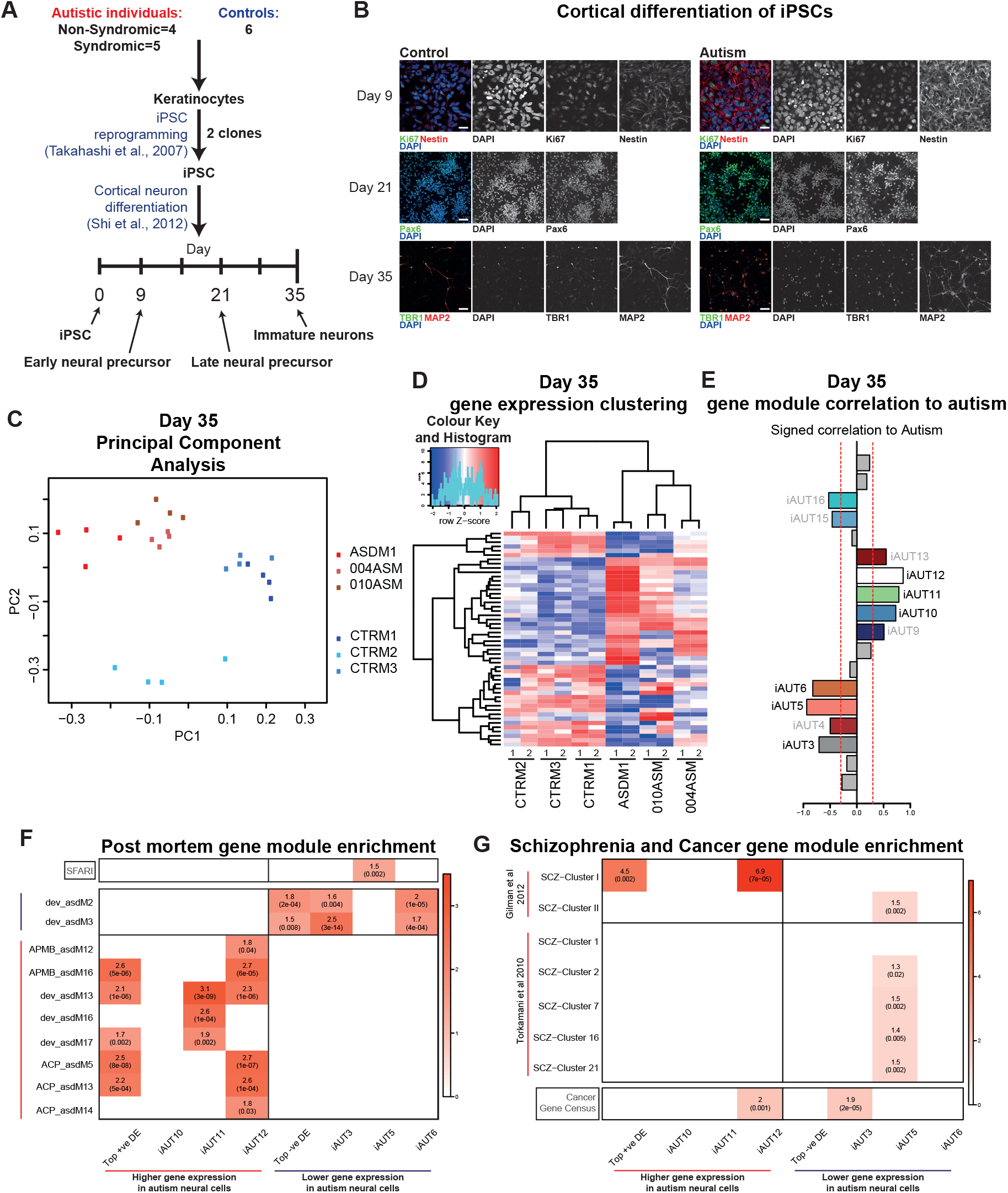
Differentiation of iPSCs into cortical lineage reveals gene expression differences between control and autism. **(A)** Study design and differentiation time points used in this study. **(B)** Differentiation of control and autism iPSCs generate precursor markers Ki67, Nestin and Pax6 and neuronal markers TBR1 and MAP2. **(C)** Principle component analysis based on gene expression counts from individual experimental replicates. **(D)** Differential gene expression and hierarchical clustering reveals significant differences between control and autism samples (biological replicates for each sample labelled 1 and 2). **(E)** WGCNA reveals 11 gene modules significantly correlated to autism (top 3 positively correlated and top 3 negatively correlated modules enrichment shown; greyed module enrichment not shown). **(F)** Gene module enrichment reveals positively correlated (red) modules are enriched in corresponding positively correlated post mortem gene modules, while negatively correlated (blue) modules are enriched in negatively correlated post mortem gene modules. **(G)** Gene modules do not show sufficient enrichment in post mortem gene modules from schizophrenia studies or cancer gene sets.

As *post-mortem* studies of adult brains have identified prenatal gene expression pathways as being altered in autism^6, 22, 23^, we were interested in determining if we could observe similar altered gene expression networks in our cohort of iPSCs. Thus, we generated day 35 neurons from three control and three non-syndromic autism-iPSCs. We chose participants with no familial history of autism or known deletions in autism-associated genes to reduce genetic bias that could drive atypical gene expression. We used an adapted bioinformatics pipeline described in^6, 23^ to analyse gene expression pathways and assess its relatedness to autism (**Supplementary Figure S2**). First, we undertook differential gene expression (DEG) and hierarchical clustering of samples based on their DEG pattern. Principle component analysis revealed significant differences between the control- and autism-iPSC neurons (**Figure 1C**), and hierarchical clustering grouped control- and autism-iPSC neurons on different branches (**Figure 1D**). Weighted gene co-expression analysis (WGCNA) revealed 11 gene modules significantly altered in autism-iPSC neurons (**Figure 1E**). The three most upregulated and three most downregulated gene modules were strongly enriched in upregulated and downregulated autism *post mortem* gene modules respectively (**Figure 1F**). These gene modules showed little to no enrichment in schizophrenia or cancer gene modules (**Figure 1G**) indicating that the gene expression patterns in our samples were autism-specific. From this we concluded that altered gene expression in adult autism brains was also found in prenatal neurons generated from iPSCs, and that gene expression patterns were specific to autism. This suggested difference between autism and typical individuals started to appear at an early stage of development.

### Marked alteration in rosette structures in autism without proliferative differences in precursor pools

Differentiation of iPSCs towards a neuronal fate first results in the generation of neuroepithelium cells. These early neuroepithelium cells become elongated and stratified and self-organise into clusters around a circular lumen known as ‘neural rosettes’^10^. These structures display apical-basal polarity similar to neural tubes^10, 24^. Critically, rosette formation and structure is thought to be key in determining cortical neurogenesis and thus generation of distinct neuronal lineages^10, 24, 25^. As our RNASeq data indicated that early neurodevelopment maybe affected in autism, we reasoned that this may be reflected by an alteration in neural rosette formation. To this end, we examined rosette formation at day 9 in control- and autism-iPSCs. As expected, all control-iPSCs robustly formed structures identifiable as neural rosettes, with an inner lumens identified by ZO-1 staining. Neural progenitor cells were observed to self-organise radially surrounding the inner lumen, typical of cells adopting an apical-basal polarity organisation (**Figure 2A**). Conversely, autism-iPSCs showed significant anomalies in lumen formation and establishment of apical-basal polarity of cells around the lumen (**Figure 2A**). We used a high content screening (HCS) approach to analyse the structure of rosettes in each line as a way to determine if there was a consistent alteration in rosette morphology between iPSC lines. All six control-iPSC line formed rosettes similar in structure with average diameters ranging between 0.066mm and 0.091mm (**Figure 2B, Supplementary Table S6**). Conversely, of the 9 autism-iPSCs, 6 formed rosettes with a smaller diameter (0.05-0.06mm); 2 did not form any rosette structures at all (026ASM and 004ASM); while 010ASM formed rosettes with diameters similar to controls (0.07mm) (**Figure 2B, Supplementary Table S6**). Autism-iPSC lines also formed more rosettes per 100 cells counted (**Figure 2C, Supplementary Table S6**). Anomalous formation of rosettes was recapitulated in 3D cortical spheroids at day 30 of differentiation (**Supplementary Figure S3A**), with fewer complete rosettes in autism-iPSC spheroids than control-iPSC spheroids (**Supplementary Figure S3B**). One explanation for these observed morphological differences could be that autism-iPSCs have altered levels of cell proliferation. Therefore, we assessed cell proliferation in day 0, 9, 21, 35 differentiated control- or autism-iPSCs. All control- and autism-iPSCs had similar rates of cell proliferation at each developmental stage examined (**Figure 2D**). Together, these data show that autism-iPSCs form anomalous rosettes independent of alterations in cell proliferation.

**Figure 2:**
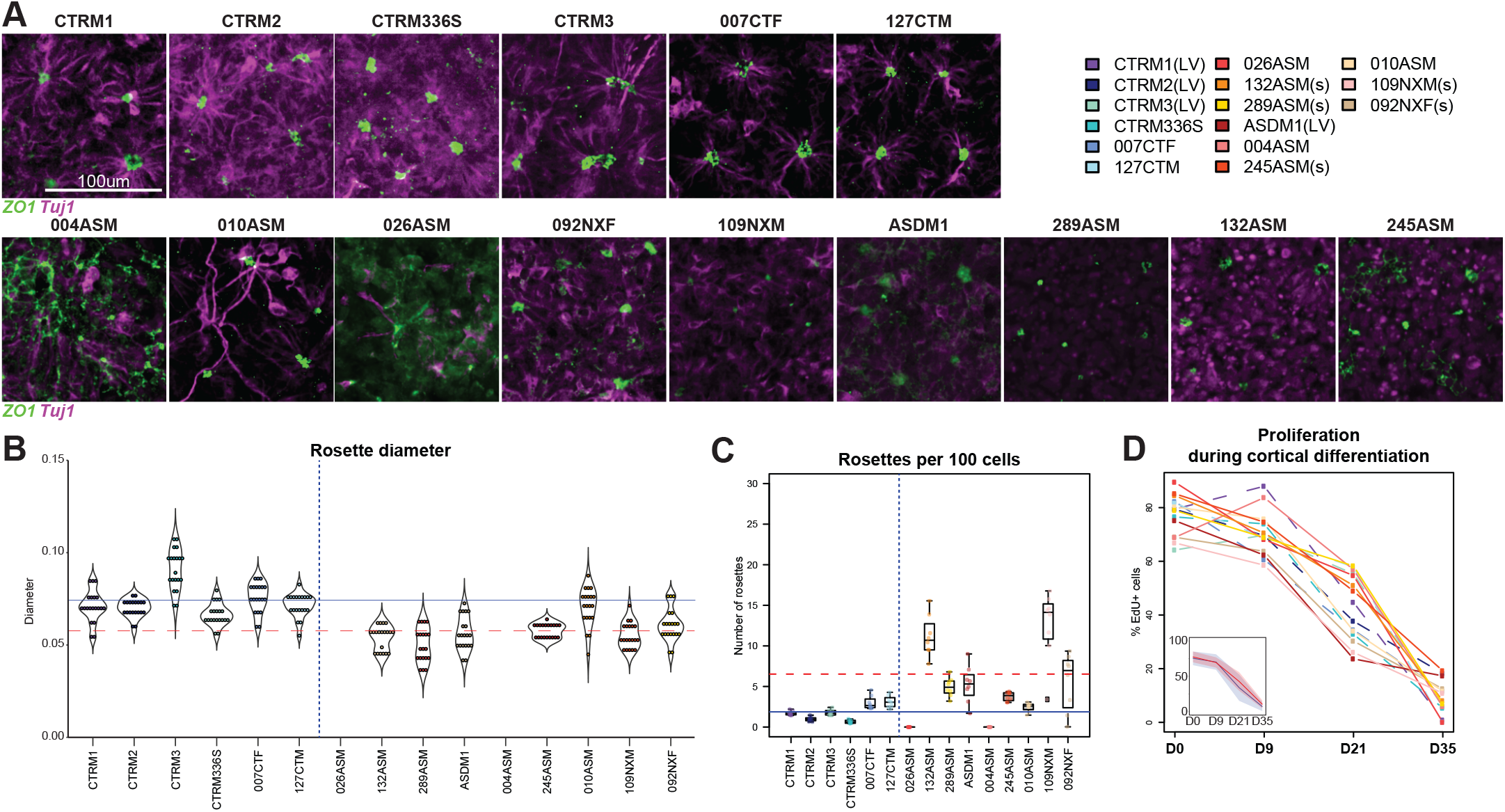
Autism iPSCs show anomalous rosette formation at day 9. **(A)** Day 9 neural rosette morphology from all participants in this study. **(B)** Rosette diameter violin plot (horizontal lines show mean rosette diameter, blue: control, red-dashed: autism). **(C)** Number of rosettes per 100 cells (horizontal lines show mean rosette number, blue: control, red-dashed: autism). **(D)** Proliferation during cortical differentiation at day 0, day 9, day 21, day 35 (dashed lines are control samples, colour key on top right corner).

### Divergence from typical development in autism occurs at a precursor cell stages during cortical differentiation

Abnormal rosette proliferation observed in autism-iPSCs could indicate premature or atypical neuronal differentiation in autism-iPSCs. To investigate this possibility, we assayed cortically differentiating iPSCs at critical stages of cortical differentiation (day 9, 21 and 35; **Figure 1A, B**) and examined the expression of fundamental cortical cell types and rate of cell type assignment at these stages using a HCS based approach. First, we asked whether control- or autism-iPSCs expressed the neuronal differentiation markers Pax6 and Tuj1 differently at early and late neural precursor cell stages (**Figure 3A**). Pax6 is a robust marker for neural precursors of cortical lineage^26^, while Tuj1 is a robust pan-neuronal and neural precursor marker^27^. Eight independent experimental replicates using 2 clones per line were assayed at every stage (**Figure 3B**). At day 9, Pax6 and Tuj1 was expressed in majority of control-iPSC cells (**Figure 3B, Table 1**). On day 21, both markers remained highly expressed (**Figure 3B**, **Table 1**). We also measure the Rate of Cell Type Assignment (dCTA) as an independent way to compare how quickly cell identity was being acquired or lost between developmental stages. In control-iPSCs, Pax6 dCTA was 13 cells/day between day 9 and day 21, while for Tuj1, dCTA=159 cells/day (**Figure 3C**). In contrast in the autism group, Pax6 and Tuj1 day 9 expression was lower than in controls (**Figure 3B, Table 1**). Assessment of cell identity acquisition in autism-iPSCs showed that Pax6 dCTA was 317 cells/day and Tuj1 dCTA=368 cells/day. These values were higher than those observed following the differentiation of control-iPSCs. However, despite this increased rate of cell identity acquisition, Pax6 and Tuj1 expression was still significantly lower in autism-iPSCs at day 21 compared to control-iPSCs (**Figure 3C, Table 1**). As expected, variability was observed throughout the differentiation protocol between experimental replicates. However, this variability was more pronounced in the autism-iPSCs. ANOVA revealed greater overall spread of data points and higher F-values in the majority of parameters assessed during differentiation of autism-iPSCs (**Supplementary Figure S4A, S4C**), while both non-syndromic and syndromic samples appeared to behave similarly (**Supplementary Figure S5A, C, D**). These data showed that control iPSC-derived precursors expressed Pax6 and Tuj1 early during differentiation, while autism-iPSCs display lower Pax6 and Tuj1 expression at the equivalent stage. Beyond this stage the rate of acquisition of Pax6 and Tuj1 was higher in autism-iPSCs, and the difference between control and autism was substantially reduced at day 21.

**Figure 3:**
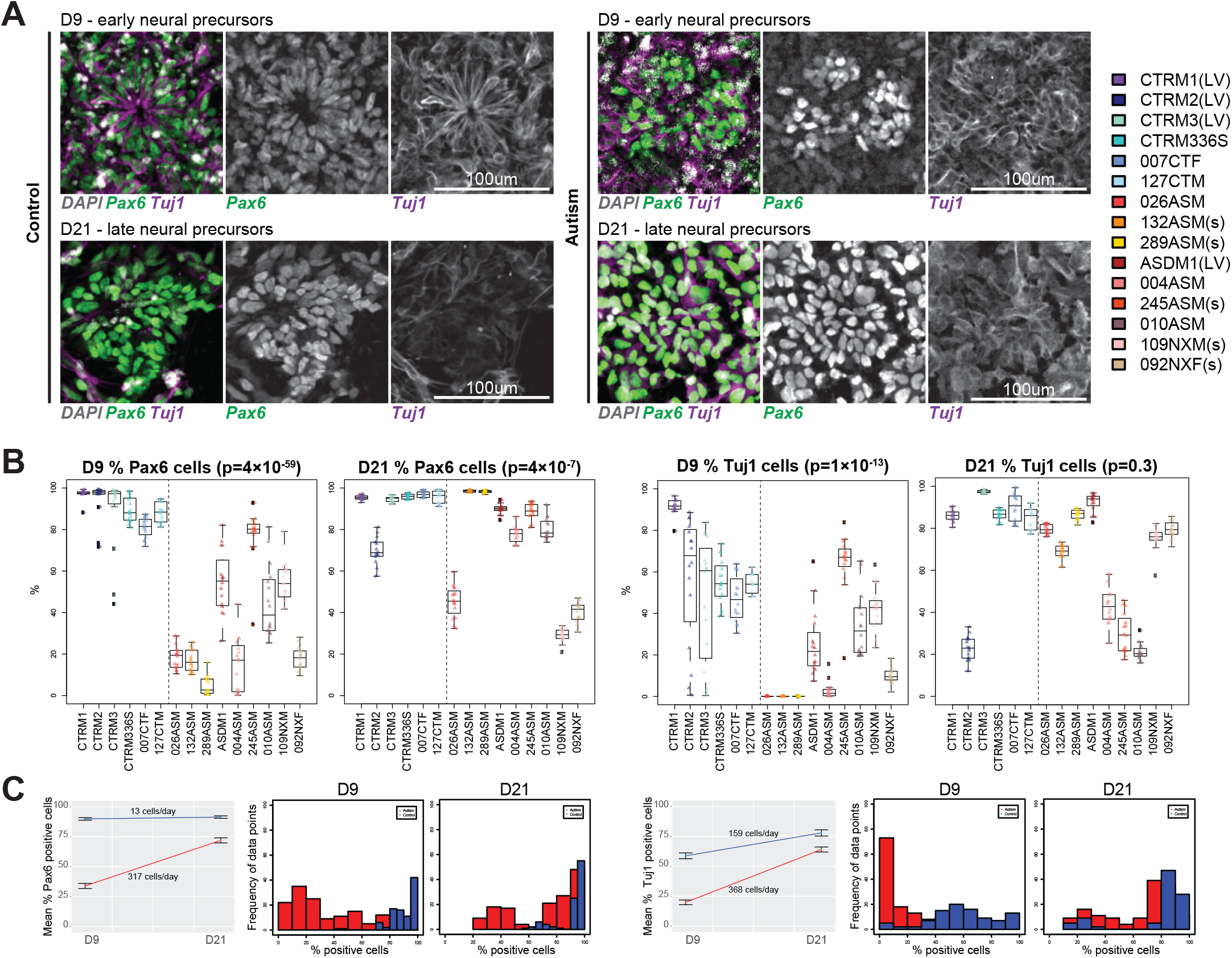
Atypical cortical differentiation of autism iPSCs. **(A)** At day 9 and day 21 precursor cell stages, both control and autism iPSCs expressed Pax6 and Tuj1. **(B)** Quantification of Pax6+ and Tuj1+ cells of individual participants (% cells positive per experimental replicate) showed significant differences between autism and control. **(C)** Mean values of % positive cells over time show significant difference between control and autism at both day 9 and day 21, as well as significant difference in rate of appearance of markers. Histogram shows normal distribution of experimental data points, and demonstrates variability between control and autism. (LV: Lentivirus reprogramming method used for generating these iPSCs; s: Participants with syndromic autism)

### Altered development of forebrain precursor lineages in autism-iPSCs independent of cell proliferation

Previous iPSC studies have linked an imbalance in GABA-glutamatergic progenitor cells and neuronal function with a macrocephaly associated cell proliferation phenotype^13, 21^. Conversely, no differences in the rate of cell proliferation throughout differentiation were observed between control- and autism-iPSCs in these studies. Thus, we were interested in establishing whether a similar imbalance in the presence of GABA-glutamatergic progenitor cells could be observed in our autism-iPSCs. To this end we investigated the development of precursors expressing Emx1, known to be expressed in dorsal forebrain (glutamatergic) neurons and precursors^28–30^, and Gad67, the rate limiting enzyme in the GABA synthesis pathway and known to be expressed in GABAergic cells^31, 32^ (**Figure 4A**). At day 9, EMX1 expression was significantly higher in control compared to autism neural precursors (**Figure 4B, C, Table 1**). At day 21, EMX1 expression in both groups appeared to remain stable, with only minor reduction in control precursors (dCTA = −41 cells/day), as opposed to a minor increase dCTA = +10 cells/day in the autism group (**Figure 4C**). At this stage, control neural progenitors expressed EMX1 significantly higher than autism neural progenitors (**Figure 4B, C, Table 1**). In day 35 immature neurons, EMX1 expression in both control and autism neurons was reduced compared to day 9 and day 21 precursors; however the reduction was significantly more acute in the autism group (dCTA = −148 cells/day in control-iPSCs vs dCTA = −254 cells/day in autism-iPSCs) (**Figure 4C**). Gad67 expression in autism- and control-iPSCs followed an opposing trajectory. At day 9, Gad67 expression was significantly higher in the control precursors, while autism precursors displayed negligible expression (**Figure 4B, C, Table 1**). At day 21, Gad67 expression was reduced in the control progenitors (dCTA = −68 cells/day), but significantly increased in autism neural progenitors (dCTA = +185 cells/day) (**Figure 4C**). Both control and autism progenitors had similar Gad67 expression at this stage (**Figure 4B, C**). However by day 35, Gad67 expression in autism neurons was higher than that in control neuron (control dCTA = −76 cells/day, autism dCTA = +176 cells/day) (**Figure 4C, D**). Similar to what we observed with Pax6 and Tuj1 expressing cells, Emx1 and Gad67 expression also showed conspicuous variability. Again, ANOVA revealed greater variability in majority of the parameters in autism lines (**Supplementary Figure S4B, S4C**), while non-syndromic and syndromic samples were similar (**Supplementary Figure S4B-D**). Lastly, we examined the expression of TBR1, a transcription factor expressed in early born excitatory neurons^10, 33^, in day 35 neurons. This revealed that differentiated control-iPSCs had higher levels of TBR1 positive cells than differentiated autism-iPSCs (**Figure 4E**). Taken together, these data showed significant differences in the determination of neuronal subtype identity of cortical lineage, in control- and autism-iPSCs. It was evident that the control- and autism-iPSCs were generating EMX1 and Gad67 expressing cells at different rates, and that there was loss of EMX1 expressing cells over time in both groups. However, at day 35 the autism-iPSCs appeared to generate greater numbers of Gad67 expressing cells over time, while in differentiated control-iPSCs, levels of both EMX1 and TBR1 remained higher than autism lines.

**Figure 4:**
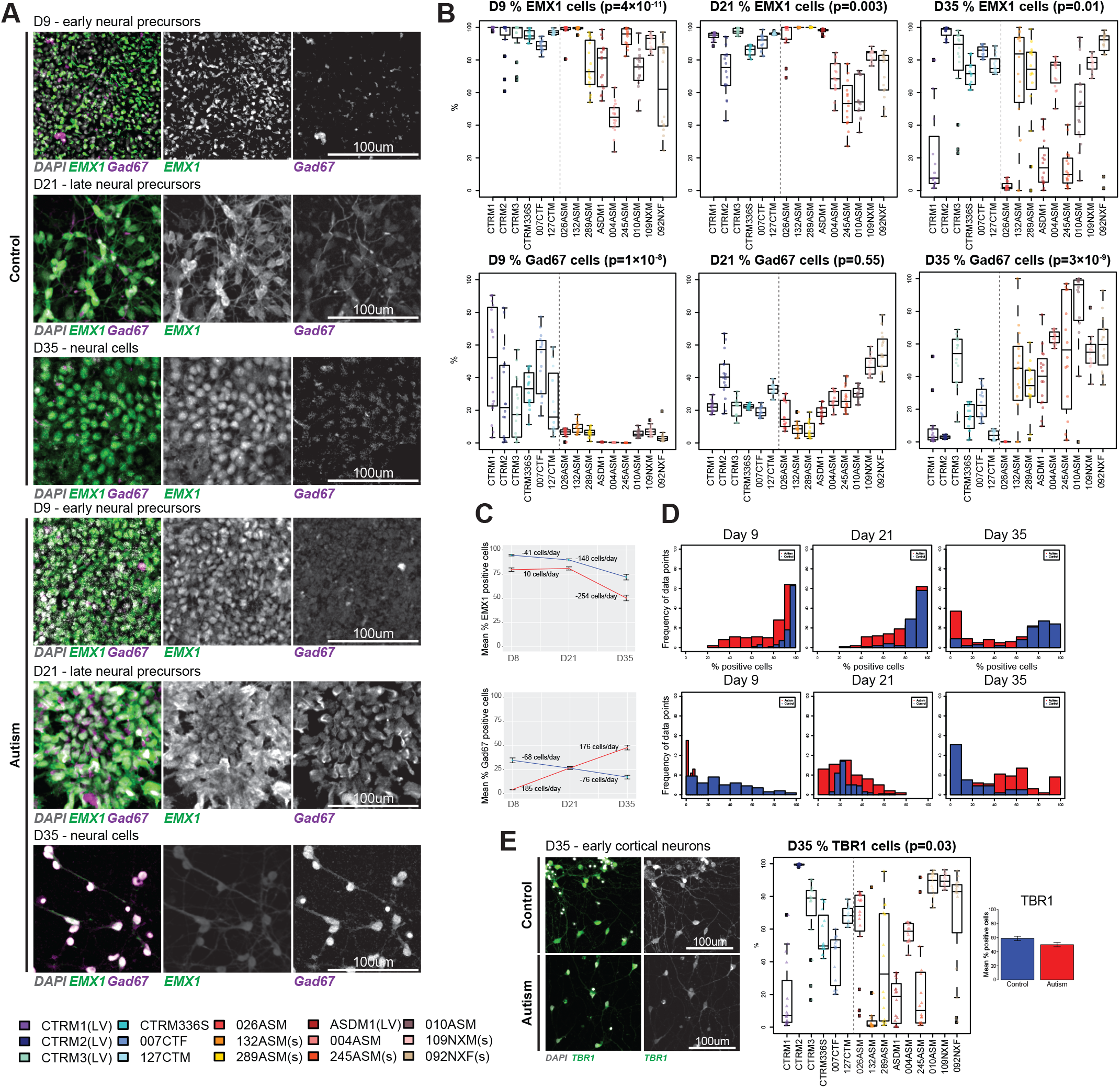
Atypical differentiation into dorsal and ventral forebrain precursors in autism. **(A)** EMX1 was expressed at day 9, day 21 and day 35 in both control and autism groups. Gad67 expression in both groups was time dependant, it decreased over time in in controls, while increased over time in autism. **(B)** Quantification of EMX1+ and Gad67+ cells (% cells positive per experimental replicate) showed significant differences between autism and control. **(C)** Mean values of % positive cells over time show significant difference between control and autism at every time point, except for Gad67 at day 21 precursor stage. **(D)** Histogram shows normal distribution of experimental data points and clear difference in distribution of data points between groups. **(E)** Control and autism iPSCs also expressed TBR1 at day 35 of cortical differentiation, and TBR1 expression was marginally higher in control vs autism. (LV: Lentivirus reprogramming method used for generating these iPSCs; s: Participants with syndromic autism)

### Generation of midbrain floorplate progenitors reveal negligible differences between control- and autism-iPSCs

The differences in cell fate acquisition observed between control- and autism-iPSCs could be due to several different factors. It could be due to genetic differences between control and autism iPSCs. Another source of variability could be attributed to stochastic fluctuations in activation of key transcription factors during differentiation, which has been reported as iPSC cells differentiate towards a cortical fate^34^. However, these differences could also be due to an inherent abnormality in the ability of our autism-iPSCs to undergo neural differentiation. Therefore, we sought to determine whether both control- and autism-iPSCs differentiated efficiently into neural progenitor cells specific for another neuronal linage; specifically mesencephalic dopamine (mDA) neurons. We chose this fate as mDAs are generated from midbrain floor plate progenitors (mFPPs) that arise from cells located on the ventral midline of the neural tube floor plate. The generation of mFPPs would, therefore, require a distinct set of factors compare to those needed for the generation of cortical precursor cells. In addition, while dysfunction in mDAs have been linked with Parkinson’s disease as well as schizophrenia, there are fewer reports of dysfunction in this population of cells in autism. Therefore, we reasoned that there would be few differences between control- and autism-iPSCs differentiating into mFPPs. This would allowing us to examine the differentiation capacity of these iPSCs. We utilized a differentiation protocol that allows for the rapid generation of a homogeneous population of mFPPs^7, 8^. After 10 days of differentiation, nearly 100% of mFPPs from both control- and autism-iPSCs were positive for LMX1A an essential transcription factor required for defining a midbrain identity^35^ (**Figure 5A, B**). No difference was observed between control- and autism-iPSCs. Similarly, expression of the transcription factor FOXA2, which positively regulates neurogenic factors in dopaminergic precursor cells^36^, did not differ between control and autism mFPPs (**Figure 5A, B**). Variability was also reduced in all the iPSC lines during midbrain differentiation (**Figure 5B**). Taken together, these data showed considerably reduced differences in midbrain lineage differentiation between control- and autism-iPSCs.

**Figure 5:**
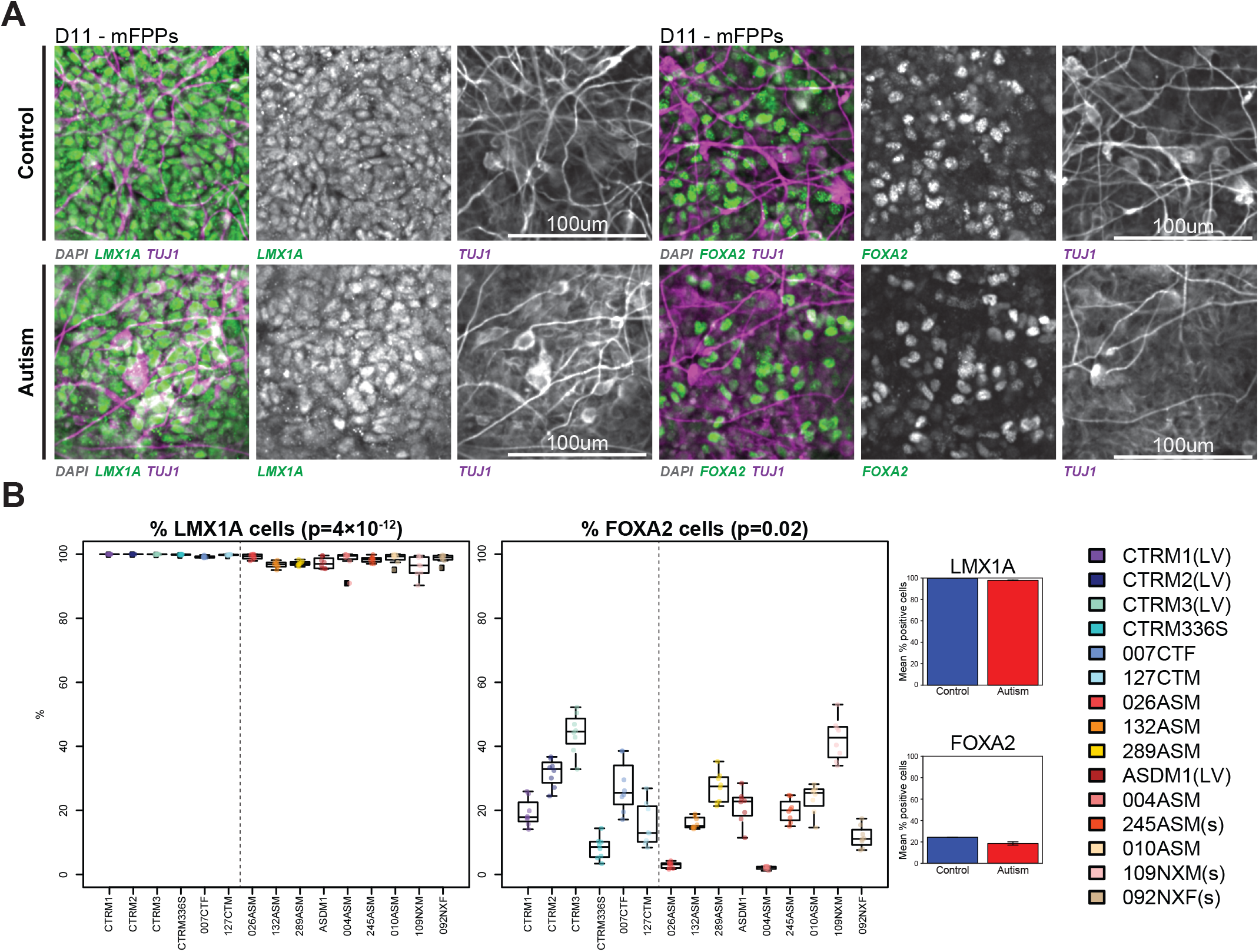
Efficient differentiation of control and autism iPSCs towards a midbrain fate. **(A)** Both control and autism iPSCs expressed LMX1A and FOXA2 when differentiated into a midbrain floor plate precursor (mFPP) cells. **(B)** Differences between control and autism iPSCs expressing LMX1A or FOXA2 was near negligible.

### Hierarchical clustering reveals sub-grouping of study participants based on cellular phenotypes alone

The findings in this study indicates that there may be a link between an autism diagnosis with atypical neurogenesis during early cortical differentiation. We have collected several cellular readouts at a number of developmental time points for each iPSC line. We reasoned that if there was a relationship between atypical cortical neurogenesis and diagnosis, that aggregating cellular phenotypes from each sample would generate a high level grouping that would distinguish between control and autistic individuals. To test this we used hierarchical clustering. This approach merges similar patterns between samples to form cluster sets, then forms bigger groups from the smaller cluster sets^37^. Being an unbiased method it can predict relatedness of samples. Data points from each participant was amalgamated into a heatmap (**Figure 6A**), and participants were ordered on the heatmap based on a mean linkage method. We then visualised the clustering in the form of an unrooted dendogram (**Figure 6B**), as participant in this study were unrelated^38^. We discovered notable relationships between samples. First, the control and autism participants grouped separately. Within the autism cluster, the participants with *NRXN1* deletions (109NXM and 092NXF) grouped on the same branch. Three syndromic autism participants (109NXM, 092NXM, 245ASM) did not group together with the non-syndromic participants. Lastly, the two autism samples 004ASM and 010ASM seemed to group on the same branch based on not only the cellular data points but also gene expression patterns as shown in **Figure 1D**. The individual patterns that emerged out of this unbiased analysis suggests that there is a potential that cellular phenotypes could reflect nature of autism diagnosis. Further studies using larger collections of deeply phenotype iPSCs as well as more detailed cellular readouts are needed in order to further understand whether such an association is robust over independent cohorts.

**Figure 6:**
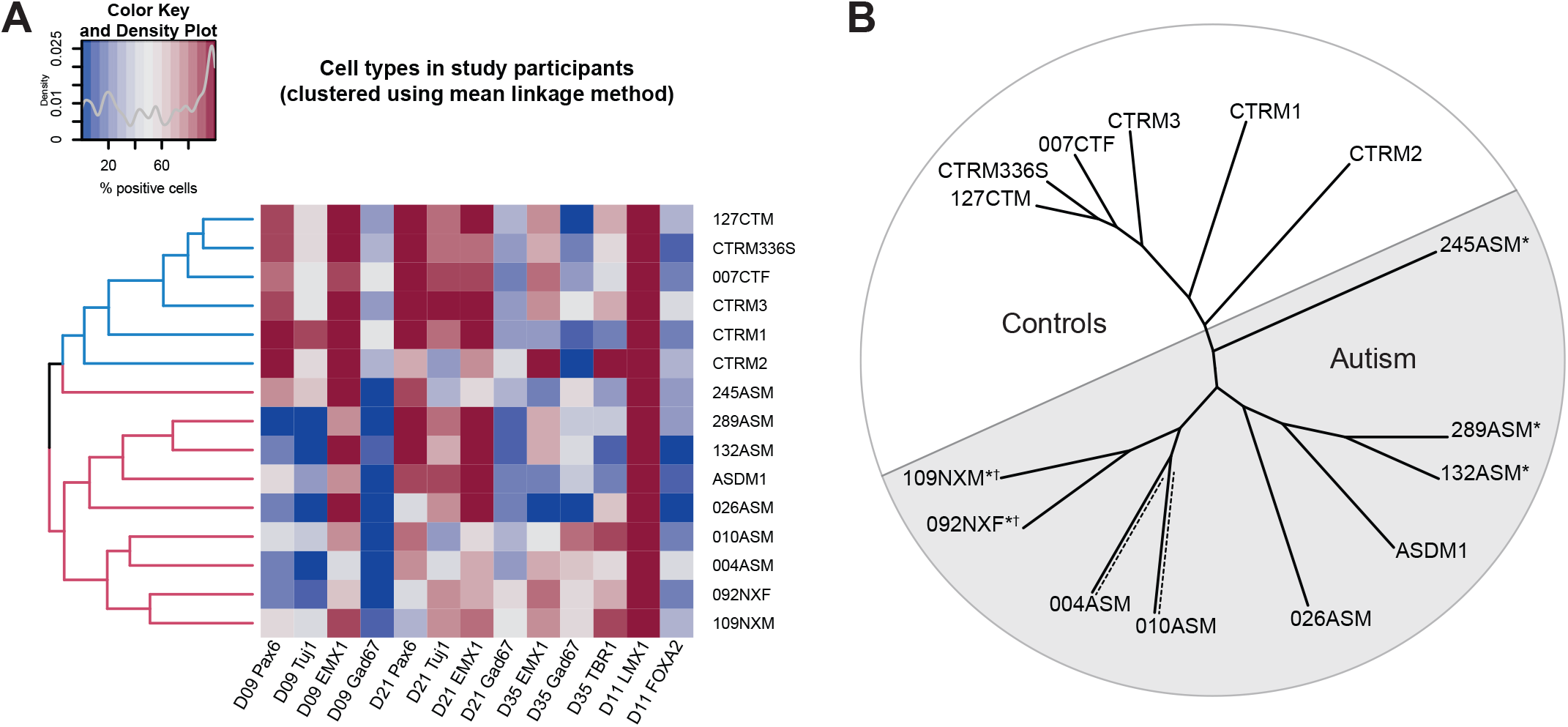
Hierarchical clustering of cellular data using mean linkage method. **(A)** All controls and autism iPSC lines were grouped based on % positive values for Pax6, Tuj1, EMX1, Gad67 at day 9, 21, 35 cortical differentiation, and LMX1A and FOXA2 at day 11 of midbrain differentiation. Controls and autism participants were grouped separately using this unsupervised learning method. **(B)** Unrooted phylogenetic tree showing relatedness of individual participants based on cellular phenotypes. Syndromic samples branched separately to non-syndromic samples*. NRXN1 deletion samples grouped together on the same branch*†. 004ASM and 010ASM which grouped on the same branch (shown with dashed lines) also grouped similarly based on gene expression data shown in Fig 1C.

## Discussion

In this study, we have investigated whether iPSCs generated from autistic individuals displayed differences during the earliest stages of cortical development. Previous studies have indicated that early development is a critical period for the emergence of phenotypes associated with autism^12–14^. However, these studies have utilized iPSCs generated from autistic individuals comorbid with macrocephaly, making it unclear whether the observed cellular effects were associated with autism or altered brain size. In this study, we have studied a collection of iPSCs generated from a heterogeneous group of autistic individuals without macrocephaly, recruited from three independent cohorts. Thus, we were able to test whether altered cellular identities occurred during differentiation of autism-iPSCs towards at cortical fate, and if this was detectable from an early developmental stage. This collection included 4 autistic individuals with uncharacterised genetic background and 5 autistic individuals with known CNVs in high risk autism loci.

First we found that autism-iPSCs generated abnormal neural rosettes, indicating an alteration in neural differentiation. Consistent with this, autism-iPSCs showed significant differences in development of early neural progenitor cells. This effect persisted at a late precursor cell stage although to a lesser degree. No differences in proliferative capacities was observed between control- and autism-iPSCs indicating this this was not the cause of altered neurogenesis in autism-iPSCs. Examination of cortical neuron subtypes revealed a divergence in the development of dorsal forebrain or excitatory precursors and ventral forebrain or inhibitory precursors from an early stage of development. Conversely, control- and autism-iPSCs demonstrated the same ability to into mFFP cells. This indicates that atypical neurogenesis predominately impacts the development of cortical linages in autism-iPSCs. Finally, based on all the temporal cortical data points acquired in this study, the participants grouped separately into controls and autism, with further unbiased branching within the autism cohort. Together, these data suggests that unique developmental differences associated with autism may be established at early prenatal stages.

We were particularly interested in modelling divergent patterns of development in the autistic cortex. We used a cortical differentiation protocol that recapitulates cortical precursor development from iPSCs, and yielded primarily excitatory cortical neurons^10^. This enabled us to study early stages of neural development, when neural rosette begin forming (day 9), equivalent to neural tube closure (approximately 4 weeks of gestation)^39^. We found marked anomalies in rosette morphology in 3 out of 9 autism-iPSCs (004ASM, 026ASM, 245ASM) resulting in either malformation or negligible neural rosette formation. In 010ASM, neural precursors were visibly dissociated from the rosette-structure, while in 092NXF, 109NXM, ASDM1, 132ASM and 289ASM cells appeared elongated and lumen formation was also affected. Further studies are required to elucidate the mechanisms responsible for the altered rosette structures and formation observed. Disruption of neural rosettes has been found to promote premature neurogenesis^40, 41^. This may explain the high rate of Pax6+ and Tuj1+ precursor generation between day 9 and day 21 in autism-iPSCs. It could also explain divergent precursor subtype assignment during early development, which we observed through opposing trajectory of Gad67 expressing cells in control- and autism-iPSCs. We noted that the appearance of Gad67+ cells in our cultures was surprising as SMAD inhibition is known to drive stem cells towards a dorsal forebrain lineage, while GABAergic neurons are known to be generated from a ventral forebrain lineage^42^. However, low numbers of GABAergic cells are known to be generated using the SMAD inhibition protocol^43, 44^, and appearance of Gad67+ cells and their dysregulation in our study may be a result of dysregulated molecular mechanisms associated with atypical precursor subtype assignment.

It is of note that in the current study the phenotypic changes occurred without the presence of proliferative differences between control- and autism-iPSCs. This suggested that cell type and structural anomalies previously reported using autism iPSCs^12, 13^ may be independent of macrocephaly associated cell proliferation alterations. Alterations in rosette formation may also contribute to the switching of precursor identity seen during development in autism-iPSCs. Further investigation into temporal precursor cell type specification will be needed to understand the mechanisms and types of cells involved. Notably, iPSC studies of non-syndromic autism remain underpowered. Nevertheless, the reports of neurodevelopmental differences between autism- and control-iPSCs are robust^13, 14, 45^. Although our cohort size would be considered inadequate for a study into non-syndromic autism, it is comparable to recent iPSC-based psychiatric studies^12–14, 46^. To achieve effect size in our study, we have used multiple clones for each iPSC-line. In addition, we utilized a HCS screening of ‘cellomic’ cell-based phenotyping approach^16, 17, 47^, recording thousands of data points from each iPSC-line.

Another consideration we faced during cellular phenotyping of iPSCs being differentiated towards a cortical fate was the high degree of variability between experimental replicates. This variability is due in part to stochastic fluctuations in transcription factor activation during cortical differentation^34, 48^. We observed that out of the 10 temporal data points recorded, 7 showed a greater degree of variability in autism-iPSCs. To rule out whether this variability was due to an iPSC-related abnormal artefact, we differentiated both control- and autism-iPSCs towards a mesencephalic fate. Following this protocol iPSCs from either control of autistic individuals behaved similarly and demonstrated reduced variability. This suggests that the variability observed in this study is specific to the cortical differentiation rather than an iPSC-related artefact. Moreover, these data indicate that alteration during early stage of development associated with autism may occur in a region specific manner.

In this study, we have used iPSCs generated from independent cohorts and from individuals with autism but without macrocephaly. Using unbiased methods, we identify that differentiation of autism-iPSCs towards a cortical but not a mesencephalic fate, results in abnormal neurogenesis characterised by premature maturation and abnormal specification of neural progenitor cells. These effects occur in the absence of altered proliferative activity between control- and autism-iPSCs. Identification of these cellular/molecular phenotypes enabled us to find common cellular pathways in a cohort having heterogeneous genetic background. In future, similarly designed studies will help identify which cellular pathways underlie these phenotypes, and may help to improve diagnosis and develop a greater understanding of the origins of autism.

## Acknowledgments

We gratefully acknowledge the participants in this study. This study was supported by grants from the European Autism Interventions (EU-AIMS) and AIMS-2-TRIALS: the Innovative Medicines Initiative Joint Undertaking under grant agreement no. 115300, resources of which are composed of financial contribution from the European Union’s Seventh Framework Programme (FP7/2007-2013) and EFPIA companies’ in kind contribution (JP, SBC, DPS, DM, GM); StemBANCC: support from the Innovative Medicines Initiative joint undertaking under grant 115439-2, whose resources are composed of financial contribution from the European Union [FP7/2007-2013] and EFPIA companies’ in-kind contribution (JP, DPS); MATRICS: the European Union’s Seventh Framework Programme (FP7-HEALTH-603016) (DPS, JP); the Wellcome Trust ISSF Grant (No. 097819) and the King’s Health Partners Research and Development Challenge Fund – a fund administered on behalf of King’s Health Partners by Guy’s and St Thomas’ Charity (Grant R130587) awarded to DPS; an Independent Investigator’s Award from the Brain and Behavior Foundation (formally National Alliance for Research on Schizophrenia and Depression (NARSAD); Grant No. 25957) to DPS, and Seed funding from Medical Research Council, UK (MR/N026063/1) awarded to DPS; the Mortimer D Sackler Foundation; the Autism Research Trust, the Chinese University of Hong Kong, and a doctoral fellowship from the Jawaharlal Nehru Memorial Trust awarded to D.A. The funding organizations had no role in the design and conduct of the study, in the collection, management, analysis and interpretation of the data, or in the preparation, review or approval of the manuscript. We are grateful to Debbie Spain and Suzanne Coghlan for participant recruitment, to Rosy Watkins, Hema Pramod, Rupert Faraway, Pooja Raval, Kate Sellers, Michael Deans and Rodrigo Rafagnin for assistance during the study, and to Aicha Massrali, Arkoprovo Paul, Bhismadev Chakrabarti, Michael Lombardo, Rick Livesey and Mark Kotter for valuable discussions. We thank the Wohl Cellular Imaging Centre (WCIC) at the IoPPN, Kings College, London for help with microscopy.

## Ethics, consent and permissions

Informed consent from participants have been taken before recruitment: Patient iPSCs for Neurodevelopmental Disorders (PiNDs) study’ (REC No 13/LO/1218).

## Consent to publish

We have obtained consent to publish from the participant to report individual patient data.

## Availability of data and materials

Sequence data have been uploaded on synapse.org. Synapse ID: syn8118403, DOI: doi:10.7303/syn8118403

## Authors’ contribution

DA, JP, JC, DPS, SBC conceived the study and wrote the first draft. VS, DHG conceived and developed bioinformatics analysis framework and analysis. DA, PN, CS, KJ responsible for sample preparation. GM was responsible for ethics application. GM, MAZ, JH, IL, DS and DM responsible for recruiting and collecting hair samples from individuals with autism and controls. All co-authors contributed to study concept, design, and writing of the manuscript. All authors read and approved the final manuscript.

