## Supplemental information for "Atypical neurogenesis in induced pluripotent stem cell (iPSC) from autistic individuals"

**Supplementary Info**

1. Extended experimental procedures
2. Supplementary Figure S1
3. Supplementary Figure S2
4. Supplementary Figure S3
5. Supplementary Figure S4
6. Supplementary table S1
7. Supplementary table S2
8. Supplementary table S3
9. Supplementary table S4
10. Supplementary table S5
11. Supplementary table S6
12. Supplementary table S7

**Extended experimental procedures**

*Study participants and neuronal differentiation*

Keratinocytes were collected from autistic participants, and typical controls without an autism diagnosis (Ethics approved, 13/LO/1218) as part of a larger European studies (EU-AIMS, STEMBANCC). All participants were Caucasian, while controls were selected if they did not have diagnosis of any psychiatric conditions. These were reprogrammed into iPSCs using previously described methods ^1, 2^. IPS cells were cultured in E8 medium (Life Technologies) with E8 supplement (Life Technologies). Cell type quantification and proliferation assays were set up on 96-well plates. 2 clones were selected from each participant, and each clone had 8 technical replicates. RNA-sequencing was performed on 2 clones from each participant, and each clone had 2 technical replicates. This design was maintained at all stages of neural differentiation recorded. Induction of neurons of cortical lineage was established using a modified dual SMADi protocol^3^. Once the cell culture reached 95% confluence, neural induction was initiated by changing the culture medium to support neural induction, neurogenesis and neuronal differentiation. A combination of N2- and B27-containing media with additives was used, henceforth called ‘neuralising medium’. N2 medium consisted of DMEM/F12 (Sigma), N2 (Gibco). B27 medium consisted of Neurobasal (Invitrogen), B27 (Gibco). Neuralising medium was supplemented with ‘dual SMADi’ 1 μM Dorsomorphin (Tocris), 500 ng/ml human Noggin-CF chimera (R&D Systems) – inhibitors of WNT pathway, BMPs and SMAD, and 10 μM SB431542 (Tocris) – inhibitor of TGFβ signaling. Noggin and dorsomorphin supresses embryonic development thereby inducing neural differentiation pathways, while SB431542 mediates loss of pluripotency. Midbrain floorplate precursors were differentiated from all participants till day 11 using previously established protocols^4, 5^. To generate cortical spheroids, iPSCs were first treated with KOSR-based hiPSC media to form embryoid bodies. Neural induction was performed using dorsomorphin and SB431542. After 6 days of neural induction, spheroids were transferred into neural maintenance media till day 30, as per established protocols^6^.

*Cortical Spheroids*

Cortical spheroids were generated using methods published by Pasca *et al.* (2015). Human iPSCs were passaged at high density (80 % confluent) as previously described. The cells were then suspended using a cell lifter in KOSR media, 10 µM ROCKi and SMAD inhibitors: 5 µM Dorsomorphin and 10 µM SB431542. The cultures were incubated in a 5% CO2 incubator and left undisturbed for 48-hours period to promote formation of embryoid bodies. Starting the second day and until the fourth day media is changed daily with fresh KOSR media supplemented with 5 µM Dorsomorphin and 10 µM SB431542. By the fifth day embryo bodies were clearly visible. At this stage, till 25th day the culture media used was B27 minus Vitamin A media plus 20 ng/mL of recombinant human EGF and recombinant human bFGF. From day 25, B27 minus Vitamin A media with 20 ng/mL of recombinant human BDNF and recombinant human NT3 was used. Media was changed every 24 hours during the first 15 days and once every 48 hours thereafter. The spheroids were harvested at day 30.

*Immunocytochemistry*

Cultures were fixed in 4% formaldehyde followed by ice-cold 100% methanol and processed for immunofluorescence staining, confocal microscopy and high throughput imaging. Secondary antibodies used for primary antibody detection were species-specific Alexa-dye conjugates (Invitrogen). We used the following primary antibodies to Ki67 (Thermo Fisher PA5-16785), Nestin (Millipore MAB5326), Pax6 (BioLegend 901301), TBR1 (Abcam ab31940), MAP2 (Abcam ab92434), Emx1 (ThermoFisher PA5-35373), Gad67 (Abcam ab26116), Tuj1 (BioLegend 801201), CD44 (R&D Systems MAB7045), LMX1A (Abcam ab139726), FOXA2 (Invitrogen 701698). Quantification was performed on the Perkin Elmer Harmony Software v4.9, which is based on the CellProfiler high throughput image analysis system^7^. Cell nuclei were first identified based on DAPI staining. For nuclear proteins, only the nuclear area was selected. For cytoplasmic protein, the area around the nucleus was selected. Thresholds of fluorescent intensity was selected after background subtraction. Threshold for each probe remained unchanged for every sample imaged. Antibodies, dilutions used and fluorescence threshold information in **Supplementary Table S5**.

Cortical spheroids were first washed in PBS and then fixed in 4% formaldehyde in 4% sucrose-PBS for 45 minutes. Following fixation, spheroids were washed in PBS and stored in sucrose 30% (weight/volume); sucrose sinking improves their preservation. After sucrose sinking, spheroids were permeabilised in 2% normal goat serum (NGS) in 0.1% Triton X100 in PBS for 60 minutes. Spheroids were incubated in a solution of permeabilization-blocking solution containing the primary antibodies for 48 hours (**Supplementary Table S5**). The samples were washed three times for three minutes in PBS and incubated with permeabilisation-blocking solution containing the secondary antibodies and HOECHST (nuclear staining) for two hours and kept in darkness to prevent bleaching of the fluorophores. Cortical spheroids were mounted as whole tissues (without sectioning), due to their small size (1-5 mm). Cortical spheroids were imaged using a Lecia SP5 confocal microscope using a 40x oil immersion objective. Imagines were acquired as Z stacks at resolution 1024x1024px and employing multiple channels: 405 (DAPI/HOECHST, blue), 488nm (green), 561nm (red) and 633nm (far-red). A line average of three was used in each channel during point scanning to prevent random background signal due to light scattering.

*EdU labelling*

IPSCs at specific stages of differentiation were labelled with EdU (5-ethynyl-2´-deoxyuridine) using the Click‑iT EdU Assay (Invitrogen). Cells were incubated with EdU for 4 hours at 37°C, then an additional 4 hours with EdU-free media. After incubation, labelled cells were fixed and prepared for detection using the Click-iT reaction cocktail. Nuclei were labelled using Hoechst 33342. Number of EdU-labelled cells were recorded as a percentage over total number of live nuclei. Imaging and analysis were done using the Opera Phenix HCS and Harmony Analysis Software.

*RNA isolation and sequencing*

RNA from 2 technical replicates was extracted using 2 clones from each participant (total: 4 samples per participant). TRIzol (Thermo Fischer) method was used, replacing chloroform with 1-bromo-3-chloropropane (BCP; Sigma). To remove genomic DNA during processing, turbo DNase (Thermo Fischer) was used. RNA concentration was quantified using Ribogreen assay (Invitrogen).

Starting with 500ng of total RNA, poly(A) containing mRNA was purified and libraries were prepared using TruSeq Stranded mRNA kit (Illumina). Unstranded libraries were constructed and underwent 50bp single ended sequencing on an Illumina HiSeq 2500 machine. To analyse iPSC mRNA-seq data, the raw reads were mapped to the human genome GRCh37.75 (UCSC version hg19) using STAR: RNA-seq aligner^8^. Aligned reads were sorted using samtools^9^, while biases were removed using Picard tools (Broad Institute). Quality control was performed using Picard tools (Broad Institute) and QoRTs ^10^. Gene expression levels were quantified using an union exon model with HTSeq ^11^, which uses uniquely aligned reads. Only the genes with >10 reads and expressed in 80% of the samples, were kept. The resulting read counts were log2 transformed and GC content, gene length, and library size normalised using the cqn package ^12^ in R.

*mRNA weighted co-expression network analysis*

Co-expression network analysis was performed using the R library, WGCNA^13^. We wanted to investigate autism-specific iPSC-neuronal culture co-expressed genes (or modules). Biweighted mid-correlations were calculated for all pairs of genes, then a signed similarity matrix was created. In the signed network, the similarity between genes reflects the sign of the correlation of their expression profiles. The signed similarity matrix was then raised to power β to emphasize strong correlations on an exponential scale. The resulting matrix (known as adjacency matrix) was then transformed into a topological overlap matrix. Since we are primarily interested in exploring co-expressed genes conserved across our cohort, we created consensus networks correlated to autism as previously published ^14, 15^. After scaling for each individual network (consensus scaling quantile = 0.2), a soft thresholding power of 14 was chosen (as it was the smallest threshold that resulted in a scale-free R­^2^ fit of 0.8) (Fig S5). The consensus network was created by using a topological overlap matrix (TOM) to calculate the component-wise minimum values for topological overlap. Using dissTOM = 1 – TOM as distance measure, genes were hierarchically clustered. Modules were then assigned using a dynamic tree-cutting algorithm (cutreeHybrid, using default parameters except deepSplit = 4, cutHeight = 0.999, minModulesize = 100, dthresh=0.1 and pamStage = FALSE).

Resulting modules of co-expressed genes were used to calculate module eigengenes (MEs; or 1^st^ principal component of the module). MEs were correlated to biological traits, in this case autism, to find disease-specific modules. Module hubs were defined by calculating module membership (kME) values which are the Pearson correlation between each gene and corresponding ME, and genes with kME < 0.7 were removed from the module. Network visualisation was done using iGraph package in R ^16^. Differentially expressed genes, and gene module assignments in **Supplementary Table S7**.

*Enrichment analysis for gene sets*

Two types of gene set enrichments were performed. For autism-correlated module enrichment, logistic regression was performed using already published gene modules ^14, 15, 17^ to control for gene length and gene expression level. A two-sided Fisher exact test with 95% confidence interval was performed for cell-type enrichment analysis using published human brain dataset ^18^.

Module genes were characterised using GO Elite (version 1.2.5) ^19^ using total expressed genes as background. GO Elite uses a Z-score approximation of hypergeometric distribution to assess term enrichment, and removes redundant GO or KEGG terms to give a concise output. 10,000 permutations were used, and required at least 10 genes to be enriched in a given pathway at a Z-score of at least 2. Only biological process and molecular function categories are reported. Pipeline schematic in **Supplementary Figure S5**.

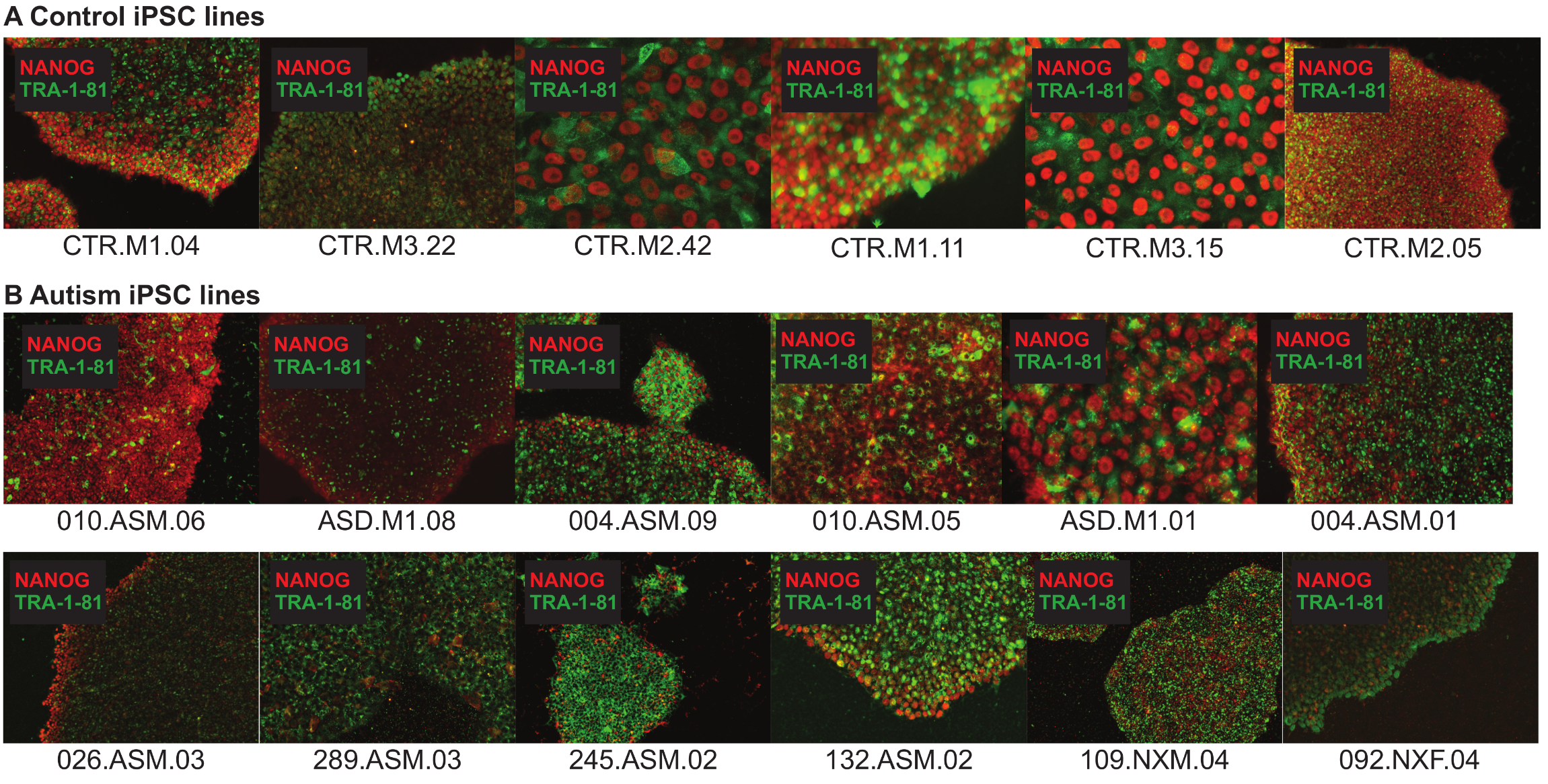
**Supplementary Figures**

**Supplementary Figure S1: Quality control images of iPSC lines.** Pluripotency of all iPSC lines were determined by positive staining for stem cell markers NANOG and TRA-1-81.

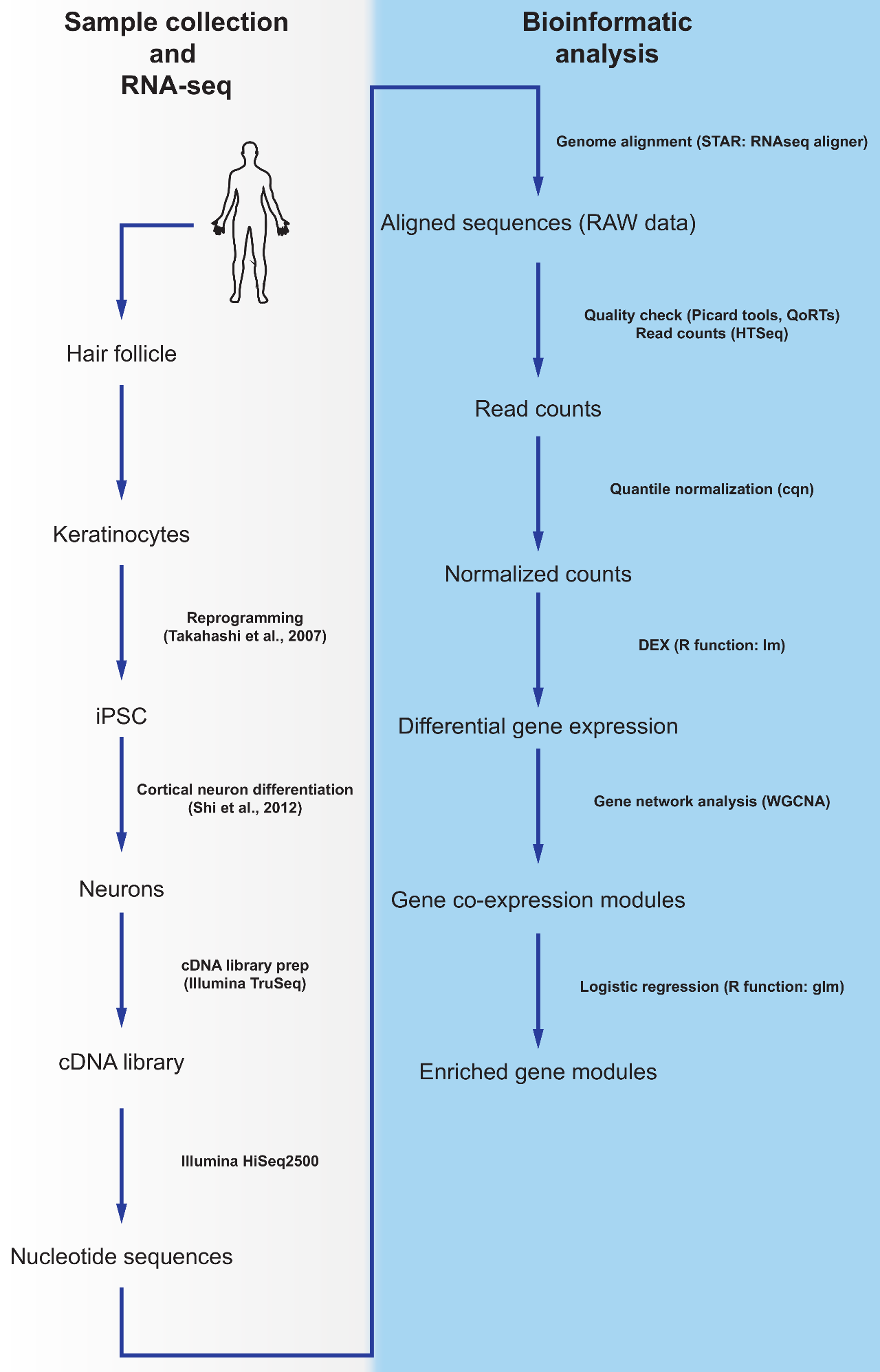

**Supplementary Figure S2: Bioinformatics pipeline.** Analysis pipeline for RNASeq.

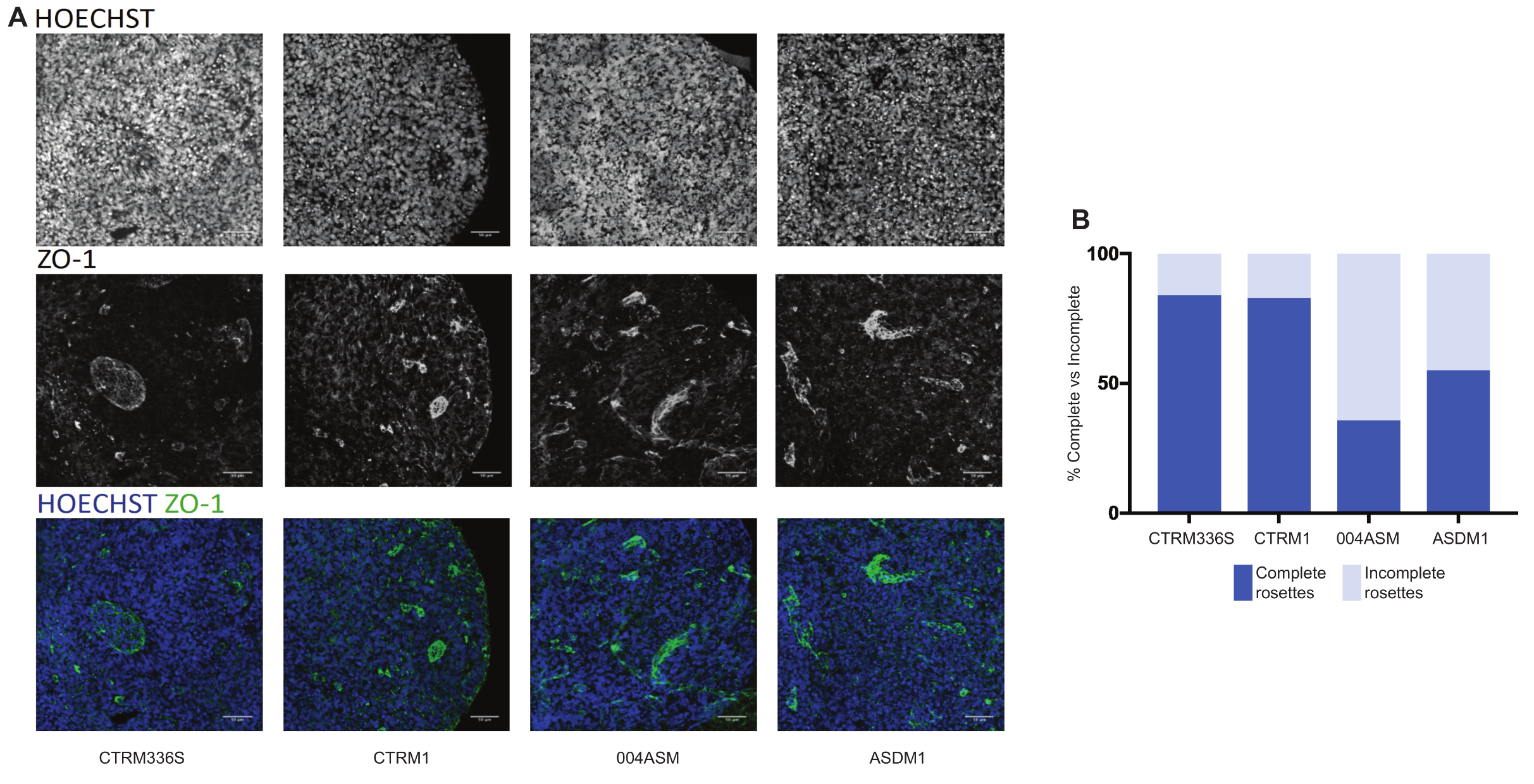

**Supplementary Figure S3: Day 7 3D spheroids from contro-l and autism-iPSCs demonstrate deficiency in neural rosette formation.** **(A)** Day 30 spheroids were immunostained for ZO1 (green) and Hoescht (Blue). **(B)** Percent complete and incomplete rosettes.

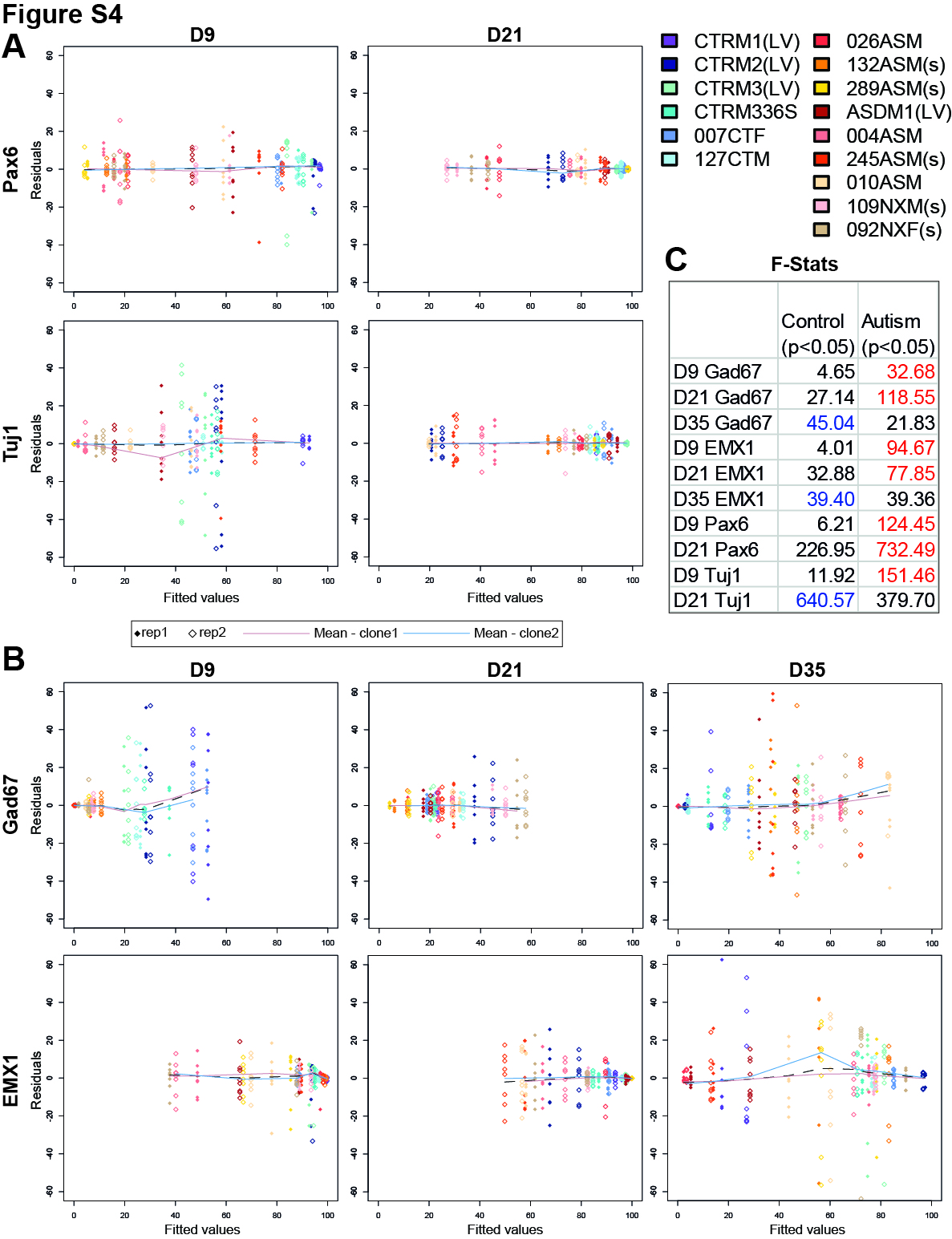

**Supplementary Figure S4: ANOVA plots to demonstrate changes in variance of data points due to clones from individual participants. (A)** Two-way ANOVA (Lines~Cells+Clones) Residuals vs Fitted values plots when observing cortical differentiation. **(B)** Two-way ANOVA (Lines~Cells+Clones) Residuals vs Fitted values plots demonstrate spread of values of individual data points across all samples when observing dorsal vs ventral forebrain differentiation. **(C)** F-values show degree of variance within the control and autism groups. All parameters measured show significant variance (p < 0.05) across both groups, and 7 out of 10 parameters show greater variance in the autism group.

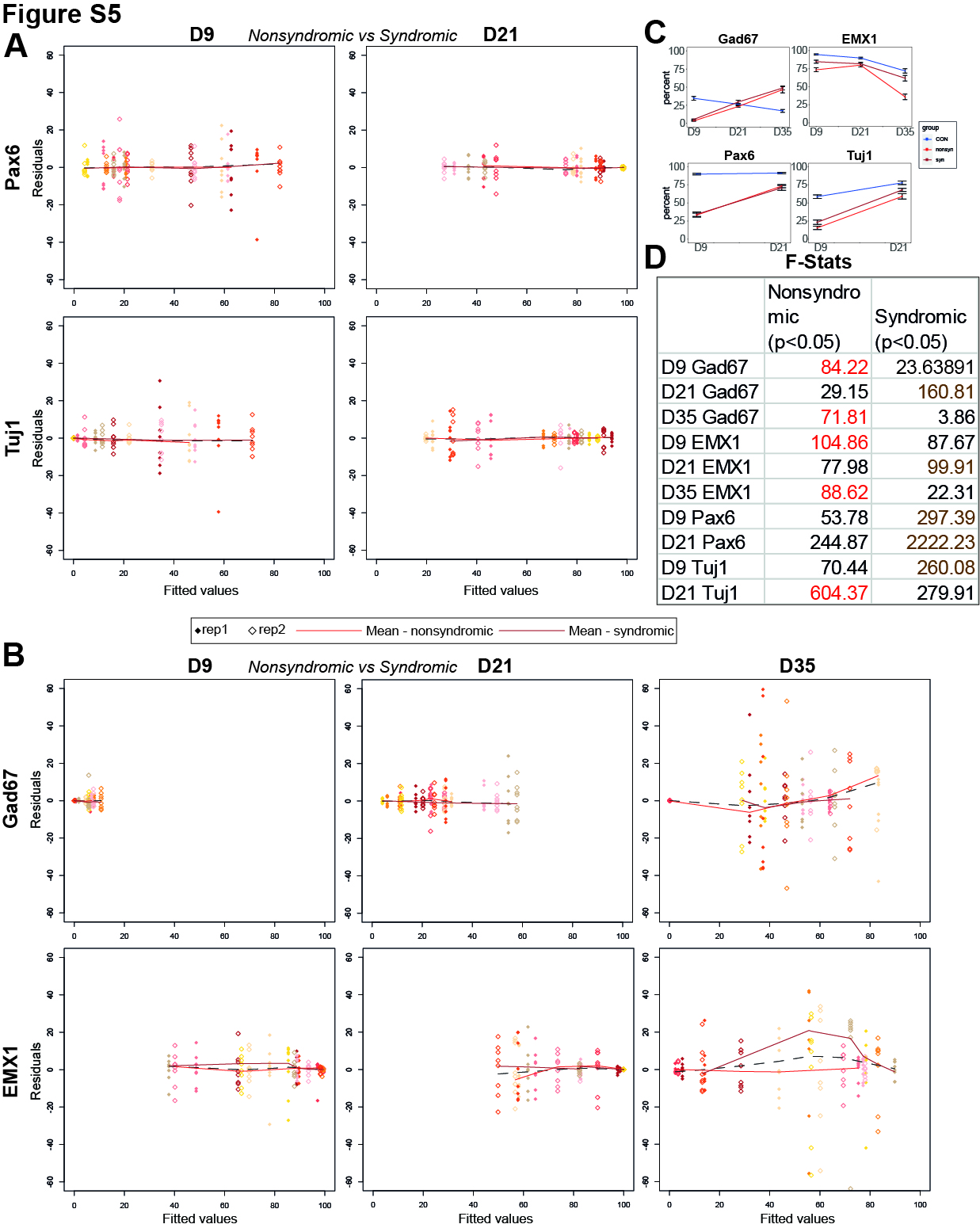

**Supplementary Figure S5: ANOVA plots to demonstrate changes in variance of data points due to type of diagnosis (non-syndromic vs syndromic) from individual participants. (A)** Two-way ANOVA (Lines~Cells+Syndromic) Residuals vs Fitted values plots when observing non-syndromic vs syndromic cortical differentiation in autistic samples.

**(B)** Two-way ANOVA (Lines~Cells+Syndromic) Residuals vs Fitted values plots demonstrate spread of values of individual data points across all autistic samples when observing non-syndromic vs syndromic dorsal vs ventral forebrain differentiation. **(C)** Mean values of % positive cells in controls, nonsyndromic and syndromic autism plotted over time. **(D)** F-values show degree of variance within the non-syndromic and syndromic autism groups. All parameters measured show significant variance (p < 0.05) across both groups, and each group shows greater variance compared to the other in equal number of parameters, 5 out of 10.

| **Table S1: General information on participants** | | | |  |  |  |  |  |  |
| --- | --- | --- | --- | --- | --- | --- | --- | --- | --- |
| Donor No. | Cohort | Ethics | iPSC ID | Autism Diagnosis | Comorbidity | Head size | Sex | Age (years) | Ethnicity |
| 1 | EU-AIMS LEAP | PINDS | ASDM1 | Yes |  | Normal | M | 34 | Caucasion |
| 2 | EU-AIMS LEAP | PINDS | 004ASM | Yes |  | Normal | M | 26 | Caucasion |
| 3 | EU-AIMS LEAP | PINDS | 010ASM | Yes | ADHD/Depression | Normal | M | 27 | Caucasion |
| 4 | EU-AIMS LEAP | PINDS | 026ASM | Yes |  | Normal | M | 16 | Caucasion |
| 5 | GOS-ICH | PINDS | 289ASM | Yes |  | Normal | M | 19 | Caucasion |
| 6 | GOS-ICH | PINDS | 132ASM | Yes | ADHD | Normal | M | 8 | Caucasion |
| 7 | BBGRE | PINDS | 245ASM | Yes |  | Normal | M | 15 | Caucasion |
| 8 | BBGRE | PINDS | 092NXF | Yes |  | Normal | F | 10 | Caucasion |
| 9 | BBGRE | PINDS | 109NXM | Yes |  | Microcephaly | M | 9 | Caucasion |
| 1 | StemBANCC | PINDS | CTRM1 | No |  | Normal | M | 58 | Caucasion |
| 2 | StemBANCC | PINDS | CTRM2 | No |  | Normal | M | 33 | Caucasion |
| 3 | StemBANCC | PINDS | CTRM3 | No |  | Normal | M | 37 | Caucasion |
| 4 | StemBANCC | PINDS | CTRM336S | No |  | Normal | M | 37 | Caucasion |
| 5 | EU-AIMS LEAP | PINDS | 127CTM | No |  | Normal | M | 56 | Caucasion |
| 6 | EU-AIMS LEAP | PINDS | 007CTF | No |  | Normal | F | 27 | Caucasion |

| **Table S2: Diagnosis details of autistic participants** | | | | |  |  |  |  |  |  |  |  |
| --- | --- | --- | --- | --- | --- | --- | --- | --- | --- | --- | --- | --- |
| Donor no. | Cohort | iPSC ID | Autism Diagnosis | ADOS |  |  |  | ADI-R |  |  |  |  |
|  |  |  |  | Social Interaction | Communication | Imagination | Stereotypic behaviour | PD_B | PD_C | PD_CS | PD_D | BDI |
| 1 | EU-AIMS LEAP | ASDM1 | Yes | 4 | 4 | 1 | 3 |  |  |  |  | 22 |
| 2 | EU-AIMS LEAP | 004ASM | Yes | 8 | 4 | 1 | 2 | 25 | 19 | NA | 4 | 3 |
| 3 | EU-AIMS LEAP | 010ASM | Yes | 5 | 5 | 1 | 2 | 3 | 3 | NA | 3 | 25 |
| 4 | EU-AIMS LEAP | 026ASM | Yes | 9 | 10 | NA | 9 | 23 | 22 | NA | 1 |  |
| 5 | GOS-ICH | 289ASM | Yes; syndromic | 3 | 3 | 0 | 1 | 9 | 5.7 | 2.3 | 1.3 |  |
| 6 | GOS-ICH | 132ASM | Yes; syndromic | 10 | 9 | NA | 9 | 15.8 | 14 | 7.5 | 5 |  |
| 7 | BBGRE* | 245ASM | Yes; syndromic |  |  |  |  |  |  |  |  |  |
| 8 | BBGRE* | 092NXF | Yes; syndromic |  |  |  |  |  |  |  |  |  |
| 9 | BBGRE* | 109NXM | Yes; syndromic |  |  |  |  |  |  |  |  |  |
| *Autism diagnosis was confirmed by referring physicians for BBGRE cohort participants. No further information available. | | | | | | | | |  |  |  |  |

| **Table S3: Genetic data of autistic participants** | | | |
| --- | --- | --- | --- |
| Donor no. | Cohort | iPSC ID | Known Deletions/Duplications/SNVs |
| 1 | EU-AIMS LEAP | ASDM1 |  |
| 2 | EU-AIMS LEAP | 004ASM |  |
| 3 | EU-AIMS LEAP | 010ASM |  |
| 4 | EU-AIMS LEAP | 026ASM |  |
| 5 | GOS-ICH | 289ASM | SNV- Chr19:41759516; C->T; AXL gene; STOP_GAINED; LoF, Synaptic transmission (Schizophrenia). CNV VOUS: 8q21.12 to q21.13 del - 8;79,886,962- 80,149,513. |
| 6 | GOS-ICH | 132ASM | CNV Clinical abnormality - 1p21.3 del - 1:96,953,361- 97,711,563 (758,201bp, DPYD, PTBP2). VOUS: 3q25.33 to q26.1 del- 3:160,629,302- 160,727,203 (97,900bp, PPM1L); 8p11.23 Dup- 8:37,124,969- 37,186,580 (61,611bp); 15q26.1 dup- 15:92,818,328- 92,875,717 (57,388bp); 17q24.1 del- 17:63,288,669- 63,375,080 (86,410bp); |
| 7 | BBGRE | 245ASM | Complete duplication of CNTN6; paternal x3 chr3:4,354,703-4,532,449 (partail dup SETMAR; complete duplication of SUMF1) |
| 8 | BBGRE | 092NXF | Paternally inherited duplication in long arms of chromosome 1 – likely to be benign – 1q21.1 (144,679,874 – 145,747,269) x3. De novo deletion of ~200kb in short arm of chromosome 2 – 2p16.3 (50,806,991 – 51,013,685) x1 |
| 9 | BBGRE | 109NXM | Maternally inherited deletion ~60kb in short arm of chromosome 2 – 2p16.3 (50,888,852 – 50,947,729) x1 |

|  | **Table S4: Reprogramming method and iPSC validation** | | |  |  |
| --- | --- | --- | --- | --- | --- |
| **Donor no.** | **iPSC ID** | **Reprogramming method** | **hiPSC validation** | **Diagnosis** | **Diagnosis type** |
| 1 | ASDM101 | Constitutive Polycistronic Lentivirus Reprogramming Kit* | ICC; cytoSNP | Autism | Nonsyndromic |
|  | ASDM108 | Constitutive Polycistronic Lentivirus Reprogramming Kit* | ICC; cytoSNP | Autism | Nonsyndromic |
| 2 | 004ASM01 | CytoTune-iPS Sendai Reprogramming Kit | ICC; cytoSNP | Autism | Nonsyndromic |
|  | 004ASM09 | CytoTune-iPS Sendai Reprogramming Kit | ICC; cytoSNP | Autism | Nonsyndromic |
| 3 | 010ASM05 | CytoTune-iPS Sendai Reprogramming Kit | ICC; cytoSNP | Autism | Nonsyndromic |
|  | 010ASM06 | CytoTune-iPS Sendai Reprogramming Kit | ICC; cytoSNP | Autism | Nonsyndromic |
| 4 | 026ASM03 | CytoTune-iPS Sendai Reprogramming Kit | ICC; cytoSNP | Autism | Nonsyndromic |
|  | 026ASM02 | CytoTune-iPS Sendai Reprogramming Kit | ICC; cytoSNP | Autism | Nonsyndromic |
| 5 | 132ASM02 | CytoTune-iPS Sendai Reprogramming Kit | ICC; cytoSNP | Autism | Nonsyndromic |
|  | 132ASM01 | CytoTune-iPS Sendai Reprogramming Kit | ICC; cytoSNP | Autism | Nonsyndromic |
| 6 | 289ASM03 | CytoTune-iPS Sendai Reprogramming Kit | ICC; cytoSNP | Autism | Nonsyndromic |
|  | 289ASM01 | CytoTune-iPS Sendai Reprogramming Kit | ICC; cytoSNP | Autism | Nonsyndromic |
| 7 | 245ASM02 | CytoTune-iPS Sendai Reprogramming Kit | ICC; cytoSNP | Autism | Syndromic |
|  | 245ASM04 | CytoTune-iPS Sendai Reprogramming Kit | ICC; cytoSNP | Autism | Syndromic |
| 8 | 109NXM04 | CytoTune-iPS Sendai Reprogramming Kit | ICC; cytoSNP | Autism | Syndromic |
|  | 109NXM03 | CytoTune-iPS Sendai Reprogramming Kit | ICC; cytoSNP | Autism | Syndromic |
| 9 | 092NXF04 | CytoTune-iPS Sendai Reprogramming Kit | ICC; cytoSNP | Autism | Syndromic |
|  | 092NXF09 | CytoTune-iPS Sendai Reprogramming Kit | ICC; cytoSNP | Autism | Syndromic |
|  | **Controls iPSCs** | |  |  |  |
| 1 | CTRM104 | Constitutive Polycistronic Lentivirus Reprogramming Kit* | ICC; cytoSNP |  |  |
|  | CTRM111 | Constitutive Polycistronic Lentivirus Reprogramming Kit* | ICC; cytoSNP |  |  |
| 2 | CTRM205 | Constitutive Polycistronic Lentivirus Reprogramming Kit* | ICC; cytoSNP |  |  |
|  | CTRM242 | Constitutive Polycistronic Lentivirus Reprogramming Kit* | ICC; cytoSNP |  |  |
| 3 | CTRM315 | Constitutive Polycistronic Lentivirus Reprogramming Kit* | ICC; cytoSNP |  |  |
|  | CTRM322 | Constitutive Polycistronic Lentivirus Reprogramming Kit* | ICC; cytoSNP |  |  |
| 4 | CTRM336S | CytoTune-iPS Sendai Reprogramming Kit | ICC; cytoSNP |  |  |
|  | CTRM337S | CytoTune-iPS Sendai Reprogramming Kit | ICC; cytoSNP |  |  |
| 5 | 127CTM04 | CytoTune-iPS Sendai Reprogramming Kit | ICC; cytoSNP |  |  |
|  | 127CTM10 | CytoTune-iPS Sendai Reprogramming Kit | ICC; cytoSNP |  |  |
| 6 | 007CTF10 | CytoTune-iPS Sendai Reprogramming Kit | ICC; cytoSNP |  |  |
|  | 007CTF01 | CytoTune-iPS Sendai Reprogramming Kit | ICC; cytoSNP |  |  |

*Note: The constitutive polycistronic lentivirus kit uses a single vector which has a negligible risk of insertional mutagenesis and viral reactivation

| **Table S5: Fluorescent thresholds for antibodies used in cell type analysis** | | | | |
| --- | --- | --- | --- | --- |
| **Antibody** | **Supplier** | **Cat. No.** | **Dilution** | **Fluorescent intensity threshold (a.u.)** |
| Pax6 | BioLegend | 901301 | 1:300 | 3500 |
| Tuj1 | BioLegend | 801201 | 1:500 | 2500 |
| Emx1 | ThermoFisher | PA5-35373 | 1:100 | 4000 |
| Gad67 | Abcam | ab26116 | 1:1000 | 600 |
| TBR1 | Abcam | ab31940 | 1:200 | 1000 |
| CD44 | R&D Systems | MAB7045 | 1:50 | 600 |
| LMX1A | Abcam | ab139726 | 1:100 | 1000 |
| FOXA2 | Invitrogen | 701698 | 1:300 | 3000 |

| **Table S6: Morphological details of Neural Rosettes** | | |  |  |  |  |
| --- | --- | --- | --- | --- | --- | --- |
| **Participant ID** | **Average diameter (mm)** | **T-test (p=1×10^-22^)** | **(mm)** | **Mean rosettes number (%)** | **T-test (p=6×10^-11^)** | **(%)** |
| CTRM1 | 0.070855781 | **Mean (control)** | 0.0742113 | 1.714263246 | **Mean (control)** | 1.87176 |
| CTRM2 | 0.070123668 |  |  | 0.980711145 |  |  |
| CTRM3 | 0.090845642 |  |  | 1.785184347 |  |  |
| CTRM336S | 0.066494935 |  |  | 0.65878026 |  |  |
| 007CTF | 0.07566226 |  |  | 2.981324875 |  |  |
| 127CTM | 0.071285499 |  |  | 3.110279078 |  |  |
| 026ASM | 0 | **Mean (autism)** | 0.04042801 | 0 | **Mean (autism)** | 6.52623 |
| 132ASM | 0.053815315 |  |  | 11.14960454 |  |  |
| 289ASM | 0.049887732 |  |  | 4.940428375 |  |  |
| ASDM1 | 0.055942962 |  |  | 5.255250823 |  |  |
| 004ASM | 0 |  |  | 0 |  |  |
| 245ASM | 0.057726738 |  |  | 3.7669185 |  |  |
| 010ASM | 0.070457893 |  |  | 2.453844441 |  |  |
| 109NXM | 0.055274512 |  |  | 12.55877341 |  |  |
| 092NXF | 0.060892681 |  |  | 5.558788098 |  |  |

**Supplementary Table S7:** Variability in data points due to clones. Data points showing significant variation due to clones highlighted in red.

|  | Clones F-stat | | | | | |
| --- | --- | --- | --- | --- | --- | --- |
|  | Combined | *p-val* | Control | *p-val* | Autism | *p-val* |
| D9 Gad67 | 0.279 | *0.598* | 0.478 | *0.4912* | 1.568 | *0.2129* |
| D21 Gad67 | 13.355 | *0.0003* | 1.554 | *0.216* | 14.505 | *0.0002* |
| D35 Gad67 | 5.435 | *0.0207* | 4.817 | *0.03* | 0.634 | *0.428* |
| D9 EMX1 | 0.269 | *0.605* | 0.025 | *0.8751* | 0.291 | *0.591* |
| D21 EMX1 | 1.635 | *0.202* | 1.239 | *0.2688* | 0.663 | *0.417* |
| D35 EMX1 | 1.095 | *0.2965* | 0.515 | *0.475* | 0.603 | *0.4388* |
| D9 Pax6 | 11.116 | *0.001* | 0.598 | *0.441* | 13.65 | *0.0003* |
| D21 Pax6 | 0.683 | *0.41* | 1.473 | *0.2283* | 0.062 | *0.8035* |
| D9 Tuj1 | 4.19 | *0.0419* | 0.496 | *0.483* | 10.69 | *0.0014* |
| D21 Tuj1 | 0.099 | *0.7538* | 0.04 | *0.842* | 0.06 | *0.8072* |
| D35 TBR1 | 0.091 | *0.763* | 1.459 | *0.231* | 1.244 | *0.267* |
